# Causal mechanisms of a healthy lifestyle intervention for patients with musculoskeletal pain who are overweight or obese

**DOI:** 10.1101/286757

**Authors:** A Williams, H Lee, SJ Kamper, KM O’Brien, J Wiggers, L Wolfenden, SL Yoong, RK Hodder, EK Robson, R Haskins, JH McAuley, CM Williams

**Author notes:** Corresponding author: Amanda Williams.

## Abstract

We assessed the causal mechanisms of a healthy lifestyle intervention for patients with chronic low back pain and knee osteoarthritis (OA), who are overweight or obese. We conducted causal mediation analyses of aggregated data from two RCTs; which included 160 patients with chronic low back pain, and 120 patients with knee OA. Participants were randomised via one central randomisation schedule, to the intervention, or usual care. The intervention consisted of brief advice and referral to a 6-month telephone-based healthy lifestyle coaching service. Participants in the back pain trial were also offered a single physiotherapy consultation. The hypothesised primary mediator was self-reported weight, and alternative mediators were diet, physical activity, and pain beliefs. Outcomes were pain intensity, disability, and quality of life (QoL). Data were analysed using causal mediation analyses with sensitivity analyses for sequential ignorability. All mediation models were specified *a priori*. The intervention had no effect on pain intensity, disability or physical QoL. The intervention significantly improved mental QoL, however, the intervention effect was not channelled via the selected mediators. The intervention did not reduce weight, or the alternative mediators (diet, physical activity, pain beliefs), and these mediators were not associated with the outcomes (with one exception; poor diet was associated with lower mental QoL). The sensitivity analyses showed that our estimates were stable across all possible levels of residual confounding. Our findings show that the intervention did not cause a meaningful change in the hypothesised mediators, and these mediators were not associated with patient outcomes.

## Background

Low back pain and knee osteoarthritis (OA) are common musculoskeletal conditions responsible for a significant global burden.^1^ In the latest Global Burden of Disease Study (2016), low back pain ranked 1^st^ and OA, for which knee OA is the highest contributor, ranked 12^th^ among all causes of years lived with disability.^1^ Consequently, these conditions cause substantial economic strain. For example, the total annual cost to Australian society was estimated at $9.2 billion (2001)^2^ for low back pain and $23.1 billion^3^ (2008) for OA.

A number of factors potentially affect the course of low back pain and knee OA. Among those commonly reported are lifestyle risk factors and erroneous pain beliefs. For example, meta-analyses have shown that being overweight or obese is associated with the persistence of low back pain^4,5^ and is an adverse prognostic factor for knee OA.^6,7^ Given their influence on weight gain, lifestyle risk factors such as poor diet and physical inactivity are also likely to indirectly influence the course of low back pain and knee OA, via weight status.^8,9^ Independently, physical inactivity is directly associated with the persistence of low back pain^10^ and poorer physical function in people with knee OA.^11^ In addition, erroneous pain beliefs are known to adversely influence outcomes from low back pain and knee OA resulting in delayed recovery and higher disability.^12,13^

Targeting lifestyle risk factors and erroneous pain beliefs are considered important aspects of treatment programs for managing chronic low back pain and knee OA.^14,15^ We conducted two randomised controlled trials (RCTs) of complex interventions targeting weight, diet, physical activity and pain beliefs, aiming to reduce pain intensity in patients with chronic low back pain^16^, and patients with knee OA,^17^ who are overweight or obese. Standard analyses of RCTs estimate whether an intervention is effective or not.^18,19^ However, these analyses cannot provide explanations for how an intervention works, or why they do not.^20^ To do so, causal mediation analysis of RCTs can be used to determine the extent to which a selected treatment target (mediator) channels the effect of the treatment onto the primary outcome.^20^ Such analyses are important to generate evidence to refine interventions, with the aim of improving their effectiveness. For example, treatment components that target effective mediators can be prioritised and strengthened in future iterations of that intervention. Conversely, mediation analyses can also explain why an intervention is ineffective. That is, by determining whether it was the intervention that failed to influence mediators, or whether the mediators were not associated with outcomes, or both.^18,21^

The underlying mechanisms of lifestyle interventions for patients with chronic low back pain and knee OA have rarely been tested.^18^ To our knowledge, only one study of a lifestyle intervention in a similar population group has investigated treatment mechanisms. Foy et al. found that in adults with knee pain and diabetes, who were overweight or obese, a reduction in weight mediated the intervention effect on disability.^22^ Given the paucity of research, the objective of this study was to test the underlying causal mechanisms of a healthy lifestyle intervention for patients with chronic low back pain or knee OA, who are overweight or obese.

## Methods

### Study design and participants

We conducted causal mediation analyses on aggregated data from two, two-arm RCTs, both part of a cohort multiple RCT.^23,24^ Full details of the methods of each trial are outlined in Williams et al.^16,23^ (ACTRN12615000478516) and O’Brien et al.^17,24^ (ACTRN12615000490572). Briefly, all patients were recruited from a waiting list for outpatient consultation with an orthopaedic specialist at the John Hunter Hospital, New South Wales (NSW), Australia. One RCT involved 160 patients with chronic non-specific low back pain,^23^ and the other, 120 patients with knee OA.^24^ All patients across both trials had a body mass index of ≥27kg/m^2^ and <40kg/m^2^ based on self-reported weight and height. Participants were randomised to both trials via one central randomisation schedule, to receive a healthy lifestyle intervention (intervention group), or remain in the cohort follow up (usual care control group), in a 1:1 ratio. The randomisation schedule was generated *a priori* by an independent investigator using SAS 9.3 through the SURVEYSELECT procedure. The pre-specified analysis plan for the current study is outlined in Lee et al. 2017.^25^

### Intervention

In both trials, participants allocated to the intervention group received brief telephone advice provided by trained telephone interviewers immediately after baseline assessment and randomisation. This advice included information about the potential benefits of weight loss and physical activity for low back pain or knee OA. Participants were then referred to the NSW Get Healthy Service (GHS) (www.gethealthynsw.com.au).^26^ The GHS is a free public health telephone-based service provided by the NSW Government to support adults to make sustained lifestyle improvements including diet, physical activity, and achieving or maintaining a healthy weight.^26^ All GHS health coaches were trained in evidence-based advice for chronic low back pain and knee OA. This training involved a 2-hour interactive workshop and information resources to guide advice for study participants.

Participants in the chronic low back pain trial were also offered a clinical consultation with the study physiotherapist. The consultation involved a clinical assessment, patient education to correct erroneous pain beliefs and behaviour change techniques to facilitate healthy lifestyle habits and weight management, informed by Self Determination Theory.^27^ Although pain beliefs were not directly targeted in the knee OA trial, we hypothesised that promotion of physical activity by the GHS service could change pain beliefs (e.g. that pain does not need to be a barrier to a physically active lifestyle).

### Control

Participants allocated to the control group continued on the usual care pathway (i.e. remained on the waiting list to have an orthopaedic consultation and could progress to consultation if scheduled) and took part in data collection during the study period. No other active intervention was provided as part of the study, however; no restrictions were placed upon the use of other health services during the study period. Control participants were informed that a new clinical service would be available in approximately 6 months involving clinical assessment and support from other services for their back pain or knee OA should they need it. No other details about the new service, or that other patients had started this service were disclosed.

### Measures

#### Mediators

The selected primary mediator was self-reported weight, in kilograms. Alternative mediators were: physical activity measured using the Active Australia Survey,^28^ which has moderate reliability (Cohen’s Kappa = 0.52)^29^ and good face and criterion validity;^30^ dietary intake measured using a short food frequency questionnaire,^31^ which has moderate reliability (Weighted Kappa range = 0.37 to 0.85)^32,33^ and criterion validity;^33^ and pain related attitudes and beliefs measured using the Survey of Pain Attitudes One-item Questionnaire, which is strongly associated with the parent questionnaire that has acceptable levels of reliability and validity.^34,35^

#### Outcomes

The primary outcomes were average self-reported pain intensity over the previous 7-days, measured using an 11-point pain Numeric Rating Scale (0=no pain, 10=worst possible pain);^36^ self-reported disability measured using the 24-item Roland-Morris Disability Questionnaire (RMDQ) in participants with chronic low back pain,^37^ and the Western Ontario and McMaster Universities Osteoarthritis Index (WOMAC)^38^ in participants with knee OA; and physical and mental quality of life (QoL) measured using the Short Form Health Survey 12 V.2.^39^ All outcomes are widely used and validated measures for these populations.^36–40^

#### Potential confounders

We identified potential confounders of the mediator-outcome effects based on theorised causal effects on the mediator and outcome variables. The selected confounders were: duration of pain (years since onset), pain intensity, disability and QoL, all measured at baseline.

### Data collection

Participant characteristics, primary and alternative mediators, outcomes and potential confounders were measured at baseline prior to random allocation by telephone interview. The primary mediator (self-reported weight) was measured 6 months after randomisation. The alternative mediators (diet, physical activity, pain beliefs) were measured 6 weeks after randomisation. The different timing of the measurement of the primary and alternative mediators was planned *a priori* to facilitate analysis via multiple mediator models (if appropriate), as per the pre-specified analysis plan outlined in Lee et al. 2017.^25^ The outcomes (pain intensity, disability, and QoL) were measured 6 months after randomisation. All mediators and outcomes were collected by a questionnaire completed via telephone by trained telephone interviewers blind to group allocation or mailed in the post as per participant preference.

### Statistical analysis

We used causal mediation analyses to analyse the data following the pre-specified analysis plan outlined in Figure 2 of Lee et al. 2017.^25^ We conducted all analyses in R (The R Foundation for Statistical Computing) using the “mediation” package.^41^

We constructed independent single mediator models for each hypothesised mediator (weight, diet, physical activity and pain beliefs) for each outcome (pain intensity, disability, physical QoL and mental QoL). Directed acyclic graphs for each model are shown in Figure 1.

**Figure 1.**
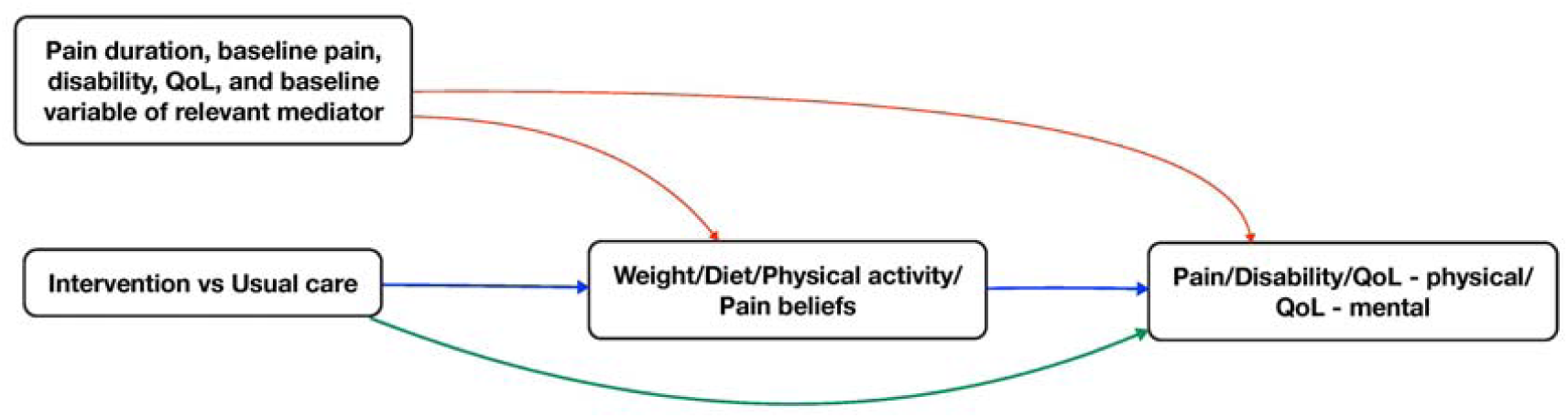
Directed acyclic graph representing a single mediator model where the intervention exerts its effect on the outcome (i.e. pain intensity/disability/QoL – physical/QoL – mental), via an indirect path (blue lines) through the mediator (i.e. weight/diet/physical activity/pain beliefs) and via a direct path (green line). Red lines represent possible effects that could induce confounding for indirect and direct effects. We assumed independence between all four mediators. Abbreviations: QoL = Quality of Life

We assumed that the intervention-mediator and intervention-outcome paths were not confounded due to random allocation of patients to intervention and control groups. However, as the mediator cannot be randomised, the mediator-outcome path is likely to be confounded. Therefore, we included theorised potential confounders (duration of pain, baseline pain intensity, disability and QoL) in the outcome regression models as covariates.

For each model, we estimated the average total effect (ATE), average causal mediation effect (ACME), average direct effect (ADE), and the proportion mediated. The ACME is the intervention effect on the outcome via the mediator; ADE is the intervention effect that is not channelled via the selected mediator; and ATE is the sum of ACME and ADE (the entire intervention effect). The proportion mediated is the fraction of ATE that is explained by ACME.

For each single mediator model, we fit two regression models: the mediator model and the outcome model. The mediator model was constructed with treatment allocation as the independent variable, and the mediator as the dependent variable. The outcome model was constructed with treatment allocation and the mediator as independent variables, the outcome as the dependent variable, and baseline measures of the mediator and the set of theorised potential confounders of the mediator-outcome path as covariates.^42^ We also included an interaction term (treatment allocation X mediator) in the outcome model to allow for a treatment-mediator interaction effect on the outcome. We used the mediate function to compute ATE, ACME, and ADE.

We planned to present the aggregate data from both trials as per our pre-specified protocol.^25^ However, given that there were some differences between the two trials, namely the clinical populations (chronic low back pain and knee OA) and the additional physiotherapy consultation exclusively delivered in the back pain trial, it seemed plausible that effects could have been moderated by trial assignment. To determine whether this was the case, we used moderated causal mediation analysis to estimate both trial-specific effects, and average effects across both trials. We decided to interpret trial-specific effects rather than averaged effects if the ACME and ADE were conditional on trial assignment.

Our mediation models were not protected against residual confounding (i.e. due to unmeasured confounders) of the mediator-outcome path. Therefore, we explored how much residual confounding would explain away the indirect effect, by using sensitivity analyses.^20^ The level of residual confounding is represented by the correlation between the residuals (error terms) from the mediator and outcome models, denoted ρ (rho). We used the medsens function to explore how varying levels of ρ (between the extremes of −1 and +1) influenced the ACME. The output provides the value of ρ at which the point estimate and CIs of the ACME includes 0 (no mediating effect). From this, we determined how strong the effect of unmeasured confounding would need to be to invalidate the estimated ACME.

#### Deviations from the pre-specified analysis plan

We made three deviations from the pre-specified analysis plan. First, the primary mediator, weight, was self-reported rather than objectively measured, this decision was made due to the availability of data. Second, we transformed the diet measure and the physical activity measure, from an ordinal and continuous scale respectively, to a binary scale to benchmark the measures against Australian Guidelines.^43,44^ A score of ‘1’ indicates meeting the guidelines (i.e. diet: 2 or more serves of fruit and 5 or more serves of vegetables per day; physical activity: participation in ≥150 minutes of moderate to vigorous physical activity per week) and ‘0’ indicates not meeting these guidelines. Third, we harmonised measures of disability (RMDQ in participants with chronic low back pain,^37^ and the WOMAC^38^ in participants with knee OA) to facilitate the interpretation of aggregate data from the two trials. We computed standardised scores for disability using the method of Van Cleave et al. 2011.^45^ These procedures are described in Text S1 in the supplementary file.

## Results

Trial assignment (chronic low back pain vs. knee OA trial) did not moderate the ACME nor ADE for all single mediator models. Thus, we present the aggregate ACME, ADE, and ATE from both trials.

### Pain intensity

The intervention had no effect on pain intensity. The intervention did not reduce the primary mediator (weight) and did not improve the alternative mediators (diet, physical activity, and pain beliefs). None of the mediators were associated with pain intensity (Table 1).

**Table 1.** Effect decomposition for each single mediator model

| Analysis | Intervention-mediator effect (path a) | Mediator-outcome effect (path b) | ATE | ADE | ACME | Proportion mediated (%) |
| --- | --- | --- | --- | --- | --- | --- |
| Pain intensity |  |  |  |  |  |  |
| Weight | 1.50 (-2.82 to 5.81) | 0.04 (-0.02 to 0.09) | 0.14 ( -0.53 to 0.84) | 0.11 ( -0.54 to 0.81) | 0.03 ( -0.12 to 0.23) | 0.04 (-2.27 to 2.46) |
| Diet | 0.79 (0.50 to 1.25)<sup>a</sup> | 0.09 (-1.41 to 1.58) | 0.11 (-0.64 to 0.82) | 0.10 (-0.65 to 0.82) | 0.01 (-0.06 to 0.08) | 0.00 (-0.71 to 1.14) |
| Physical activity | 1.11 (0.77 to 1.60)<sup>a</sup> | -0.33 (-1.62 to 0.95) | 0.13 (-0.58 to 0.85) | 0.12 (-0.58 to 0.84) | 0.01 (-0.08 to 0.12) | 0.00 (-1.20 to 1.31) |
| Pain beliefs | 0.52 (-0.71 to 1.74) | 0.05 (-0.05 to 0.14) | 0.13 (-0.62 to 0.84) | 0.12 (-0.65 to 0.82) | 0.03 (-0.05 to 0.13) | 0.01 (-1.30 to 2.18) |
| Disability |  |  |  |  |  |  |
| Weight | 1.50 (-2.82 to 5.81) | 0.01 (-0.02 to 0.03) | 0.12 (-0.14 to 0.37) | 0.12 (-0.13 to 0.34) | 0.00 (-0.08 to 0.10) | 0.03 (-2.23 to 1.85) |
| Diet | 0.79 (0.50 to 1.25)<sup>a</sup> | 0.13 (-0.31 to 0.56) | 0.13 (-0.10 to 0.37) | 0.15 (-0.09 to 0.38) | -0.02 (-0.08 to 0.02) | -0.05 (-2.08 to 1.25) |
| Physical activity | 1.11 (0.77 to 1.60)<sup>a</sup> | 0.08 (-0.34 to 0.50) | 0.14 (-0.10 to 0.38) | 0.14 (-0.10 to 0.37) | 0.00 (-0.02 to 0.03) | 0.00 (-0.61 to 0.45) |
| Pain beliefs | 0.52 (-0.71 to 1.74) | 0.01 (-0.02 to 0.04) | 0.13 (-0.10 to 0.35) | 0.12 (-0.10 to 0.34) | 0.01 (-0.02 to 0.05) | 0.03 (-0.84 to 1.01) |
| QoL - Physical |  |  |  |  |  |  |
| Weight | 1.50 (-2.82 to 5.81) | -0.18 (-0.40 to 0.04) | -2.05 (-4.75 to 0.54) | -1.88 (-4.45 to 0.60) | -0.16 (-1.15 to 0.59) | 0.05 (-0.99 to 0.96) |
| Diet | 0.79 (0.50 to 1.25)<sup>a</sup> | 1.45 (-2.99 to 5.90) | -1.24 (-3.78 to 1.30) | -1.22 (-3.79 to 1.35) | -0.02 (-3.79 to 1.35) | 0.01 (-1.17 to 1.06) |
| Physical activity | 1.11 (0.77 to 1.60)<sup>a</sup> | 2.98 (-1.42 to 7.38) | -1.22 (-3.95 to 1.27) | -1.29 (-4.09 to 1.18) | 0.07 (-0.32 to 0.55) | -0.01 (-1.63 to 1.62) |
| Pain beliefs | 0.52 (-0.71 to 1.74) | 0.06 (-0.25 to 0.38) | -1.21 (-4.12 to 1.53) | -1.10 (-3.85 to 1.52) | -0.11 (-0.60 to 0.20) | 0.05 (-0.92 to 0.84) |
| QoL - Mental |  |  |  |  |  |  |
| Weight | 1.50 (-2.82 to 5.81) | -0.11 (-0.39 to 0.17) | 3.80 (0.48 to 7.16)* | 3.90 (0.70 to 7.02)* | -0.10 (-1.29 to 0.92) | -0.01 (-0.96 to 0.29) |
| Diet | 0.79 (0.50 to 1.25)<sup>a</sup> | -6.40 (-12.06 to -0.74)* | 4.73 (1.40 to 8.10)* | 4.62 (1.32 to 7.97)* | 0.10 (-0.53 to 0.93) | 0.03 (-0.16 to 0.29) |
| Physical activity | 1.11 (0.77 to 1.60)<sup>a</sup> | 1.78 (-4.01 to 7.58) | 4.70 (1.34 to 8.00)* | 4.69 (1.31 to 7.96)* | 0.01 (-0.37 to 0.40) | 0.00 (-0.10 to 0.11) |
| Pain beliefs | 0.52 (-0.71 to 1.74) | -0.23 (-0.64 to 0.18) | 4.92 (1.61 to 8.30)* | 5.04 (1.71 to 8.36)* | -0.12 (-0.77 to 0.30) | -0.02 (-0.20 to 0.06) |

Abbreviations: ATE = average total effect, ADE = average direct effect, ACME = average causal mediation effect, QoL = Quality of Life
Adjusted coefficients ( $\beta$ ) with their 95% confidence intervals unless otherwise stated.
<sup>a</sup>Binary model presented as an odds ratio
\*p= <0.05

### Disability

The intervention had no effect on disability. The intervention did not reduce the primary mediator (weight) and did not improve the alternative mediators (diet, physical activity, pain beliefs), and none of the mediators were associated with disability.

### Physical QoL

The intervention had no effect on physical QoL. The intervention did not reduce the primary mediator (weight) and did not improve the alternative mediators (diet, physical activity, and pain beliefs), and none of the mediators were associated with physical QoL.

### Mental QoL

The intervention significantly improved mental QoL, however, the intervention effect was not channelled via the selected mediators (Table 1). The intervention did not reduce the primary mediator (weight), and weight was not associated with mental QoL. The intervention did not improve the alternative mediators (diet, physical activity, and pain beliefs); and physical activity and pain beliefs were not associated with mental QoL. Diet was negatively associated with mental QoL (i.e. meeting the dietary guidelines for serves of fruits and vegetables per day was associated with poorer mental QoL).

### Sensitivity analyses

The sensitivity analyses showed that our estimated ACME’s were stable across all possible levels of residual confounding. The sensitivity plots for each model are reported in Figure S1 in the supplementary file.

### Multiple mediator models

As per the pre-specified analysis plan,^25^ we did not conduct multiple mediator models because the intervention did not reduce weight (primary mediator).

## Discussion

### Key findings

Our findings showed that the healthy lifestyle intervention did not improve pain intensity, disability or physical QoL in patients with chronic low back pain or knee OA. The intervention did improve mental QoL, however, the intervention effect was not channelled via the selected mediators. The intervention did not cause a meaningful change in the hypothesised primary and alternative mediators, and these mediators were not associated with the selected outcomes.

Previous studies demonstrate that interventions have successfully improved weight, diet, physical activity and pain beliefs in patients with low back pain and knee OA.^46–48^ For example, Messier and colleagues report that a 6-month diet and exercise intervention led to a mean weight loss of 8.5kg in participants with knee OA.^48^ However, most of these trials evaluated intensive face-to-face consultations and none were delivered using telephone health coaching. This difference in the mode of delivery might explain why our intervention did not exert an effect on the hypothesised mediators, whereas interventions in previous studies did. Although telephone interventions are effective in reducing weight and the behavioural determinants of weight (diet and physical activity) for the general population,^49,50^ their effectiveness for patients with chronic low back pain and knee OA have not been established.^51,52^ The telephone-based intervention used in our study was not effective in reducing self-reported weight, improving diet or physical activity, or changing erroneous pain beliefs in these patient groups.

Meta-analyses of observational cohort studies suggest that the hypothesised mediators are associated with patient outcomes.^4–6,10,13,53^ Although these meta-analyses report adjusted estimates, they did not consider the effects of unmeasured or residual confounding.^54–56^ Therefore, it is possible that these estimates were influenced by confounding bias. In our study, the ACME was stable across all possible levels of residual confounding, and we found no association between the majority of the hypothesised mediators and outcomes of pain intensity, disability, and QoL. However, in our study the absence of any association could be a result of lack of variability in the mediator scores, which may be a function of low variability in those measures in the sample at baseline and ineffectiveness of the intervention.

To our knowledge only one previous study of a lifestyle intervention in a similar population has undertaken causal mediation analyses. Foy et al. found that in adults with knee pain and diabetes who were overweight or obese, reduction in weight explained 98% of the intervention effect on disability.^22^ Conversely, we did not detect a mediating effect through weight loss. The difference in results may be because Foy et al. included patients with concomitant diabetes, which could have moderated the indirect effect. Furthermore, Foy et al. used an objective measure of weight, which may have increased the reliability and/or validity, compared to our self-reported measure. Lastly, Foy et al. did not undertake a sensitivity analysis to determine the impact of residual confounding on the mediator-outcome path, thus their estimate of the indirect effect through weight could be confounded.

Other studies suggest that improving lifestyle risk factors or changing pain beliefs positively affects patient outcomes in these patient groups.^51,57,58^ However, in the absence of causal mediation analyses, these studies can only assume that the intervention worked through hypothesised treatment targets. Without strong evidence for mediation through these targets, it remains possible that intervention acted via alternative mechanisms. Despite this uncertainty, trials without mediation analyses have informed clinical practice guidelines for chronic low back pain and knee OA. For example, for knee OA, weight loss is strongly recommended.^14^ Likewise, for chronic low back pain advice and education to correct erroneous pain beliefs is advised.^15^ Such guidelines should be better informed through robust evidence of treatment mechanisms. Collectively, the evidence to date does not convincingly demonstrate that overweight or obesity, poor diet, low levels of physical activity and erroneous pain beliefs are the appropriate mechanisms that should be targeted to improve pain intensity, disability, and QoL in patients with chronic low back pain or knee OA.

### Limitations

The hypothesised mediators in this study were measured using self-reported questionnaires. Objective measures may increase the reliability and validity of the measurement of the hypothesised mediators. The mediators, diet and physical activity, were transformed from an ordinal and continuous scale respectively, to a binary scale to allow interpretation against the existing national guidelines. This may have reduced the responsiveness of these measures. We made three deviations from the published protocol. Although we transparently disclosed these deviations, they could have introduced bias.

### Implications for future research

Although clinical guidelines advocate focusing on lifestyle risk factors and erroneous pain beliefs in patients with chronic low back pain or knee OA, there is uncertainty about whether they are causes of pain intensity, disability, and poor QoL. Future RCTs targeting lifestyle risk factors or erroneous pain beliefs in patients with chronic low back pain and knee OA should undertake mediation analyses to understand if the intervention changed the intended targets and if the targets were causally associated with the selected outcomes. To provide more convincing evidence, objective measures should be used when possible and sensitivity analyses assessing the effects of residual confounding should be undertaken.

### Clinical implications

Our study found that the healthy lifestyle intervention delivered primarily using the telephone did not change the intended targets of weight, diet, physical activity and pain beliefs. Other studies suggest that a more intensive lifestyle intervention delivered face-to-face might change these targets. Currently, we cannot recommend that a lifestyle intervention delivered by telephone is preferable over face-to-face for patients with chronic low back pain and knee OA. As it remains unclear whether the hypothesised mediators in this study are causes of pain, disability and poor QoL in patients with chronic low back pain or knee OA, it is difficult to provide clinical guidance regarding prioritisation of these mediators. However, targeting these mediators, in particular, the lifestyle risk factors, may offer other health benefits such as improved cardiovascular disease risk,^59^ particularly for overweight or obese patients.

## Conclusions

This study aimed to test the underlying causal mechanisms of a healthy lifestyle intervention for patients with chronic low back pain or knee OA who are overweight or obese. Our findings show that the intervention did not improve pain intensity, disability and physical QoL in participants with chronic low back pain and knee OA. The intervention did improve mental QoL, however, the intervention effect was not channelled via the selected mediators. The intervention did not cause a meaningful change in the hypothesised mediators, and these mediators were not associated with patient outcomes.

