## Supplementary Materials for "Causal mechanisms of a healthy lifestyle intervention for patients with musculoskeletal pain who are overweight or obese"

### Supplementary file

**Text S1:** Procedure for standardization of disability scores

The following steps were followed:

1. Transformation of raw scale scores to 0 – 100

Transformed score= $\frac{actual\ raw\ score - lowest\ possible\ raw\ score}{possible\ raw\ score\ range}\ x\ 100$

2. Calculating standard scores

Standard score= $\frac{X - \bar{X}}{standard\ deviation}$

**Figure S1.** Sensitivity analysis plots for each single mediator model with pain (1.1), disability (1.2), QoL-physical (1.3), QoL-mental (1.4) as the outcome and weight (A), diet (B), physical activity (C) or pain beliefs (D) as the mediator for the usual care control (left panel) and intervention (right panel), respectively. The correlation between the error terms in the mediator and outcome regression models (ρ) is plotted against the average causal mediation effect (ACME). The estimated ACME (assuming sequential ignorability) is the dashed line and the 95% confidence intervals are represented by the shaded regions.

**Figure1.1A**

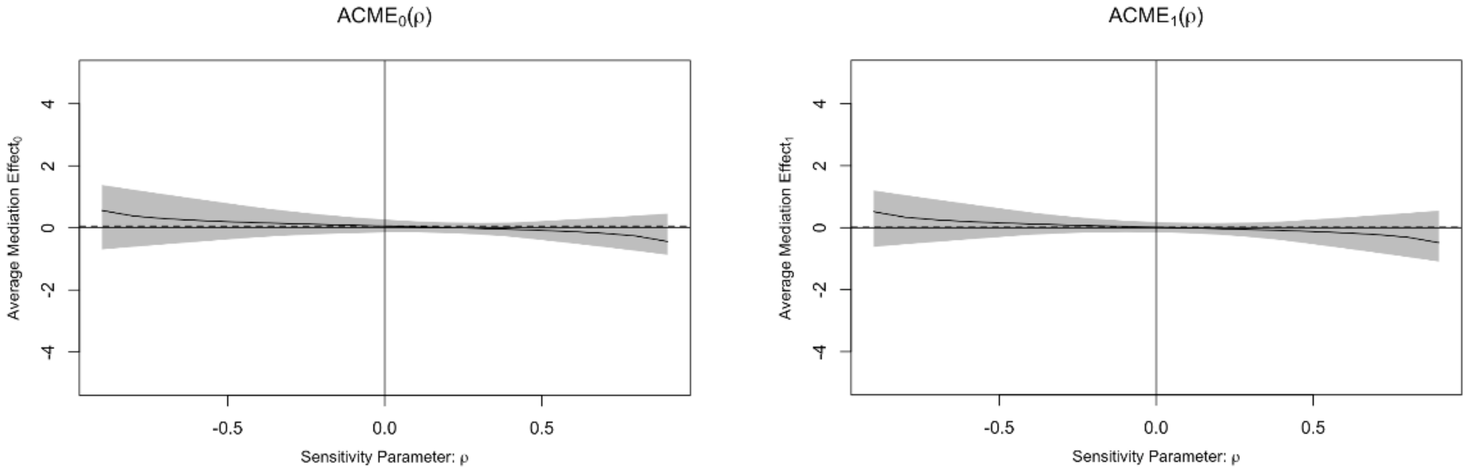


**Figure 1.1B**

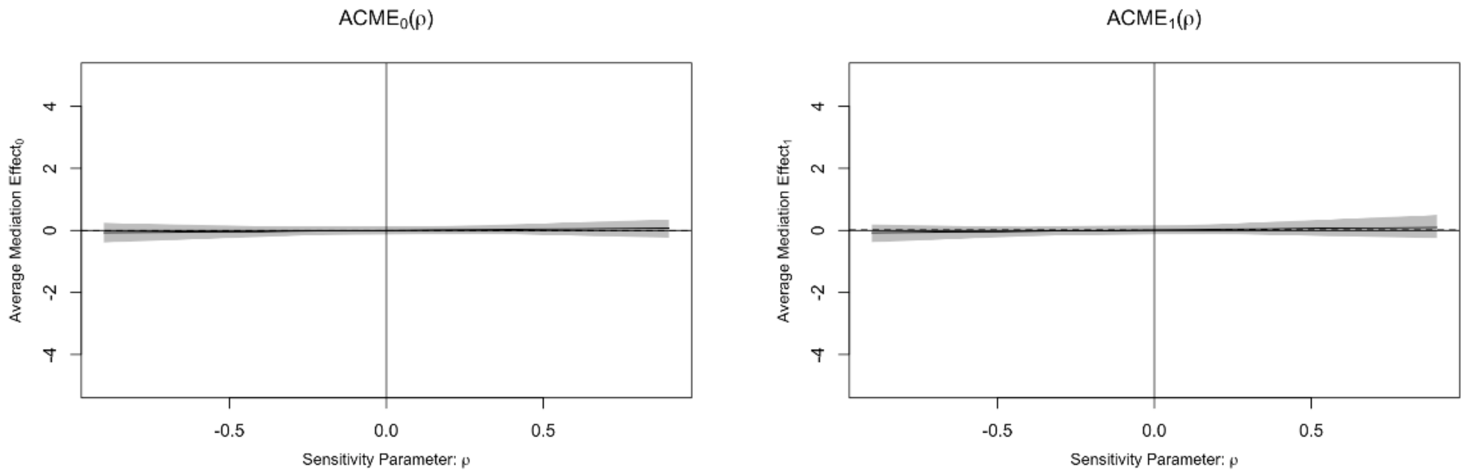

**Figure 1.1C**

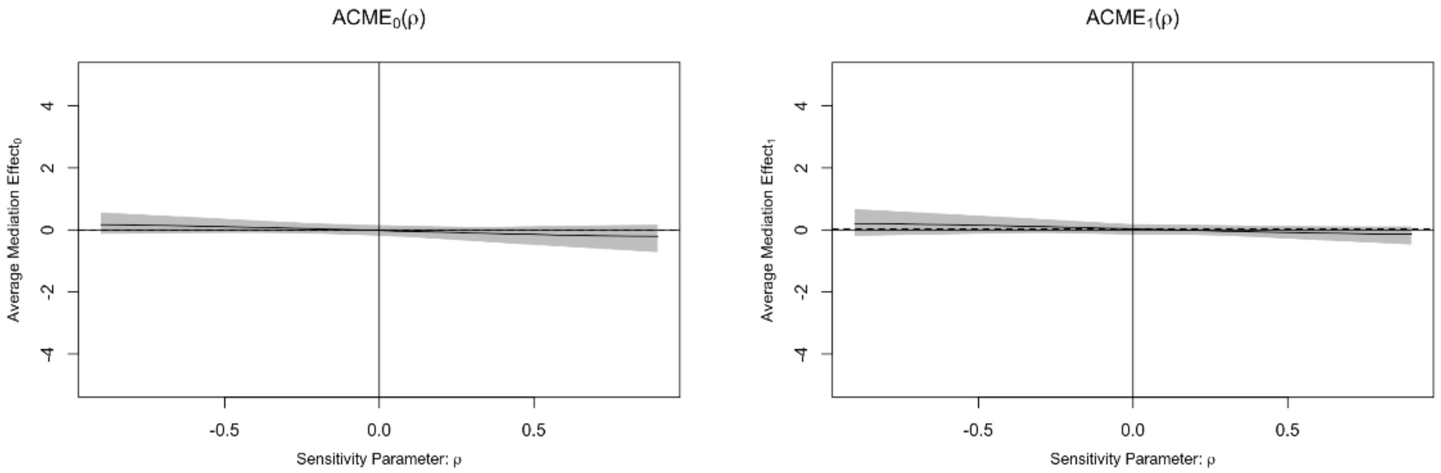


**Figure 1.1D**

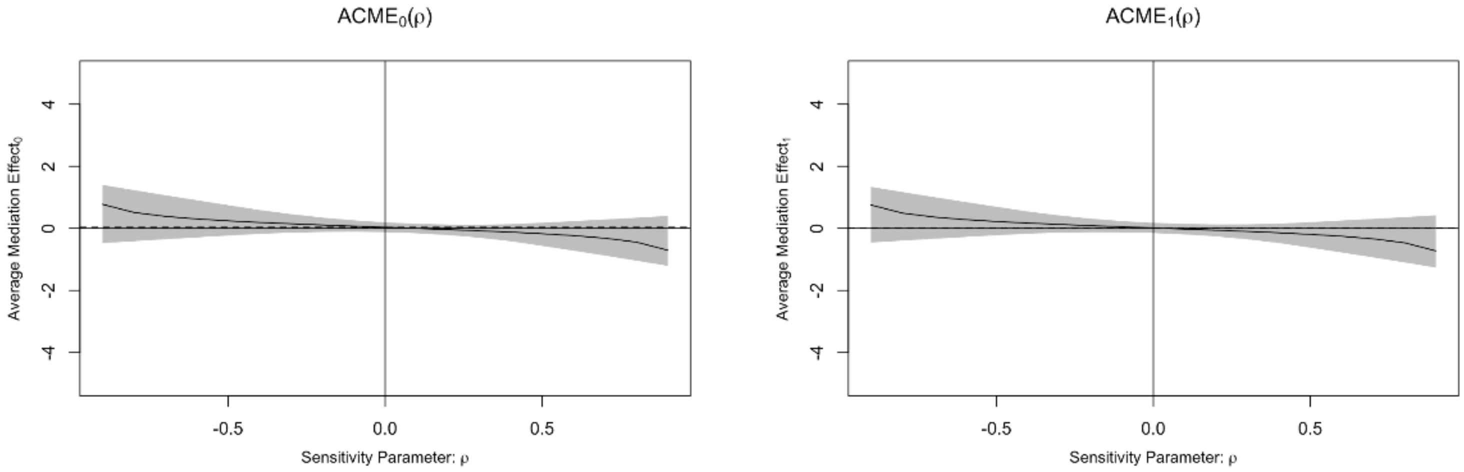


**Figure 1.2A**

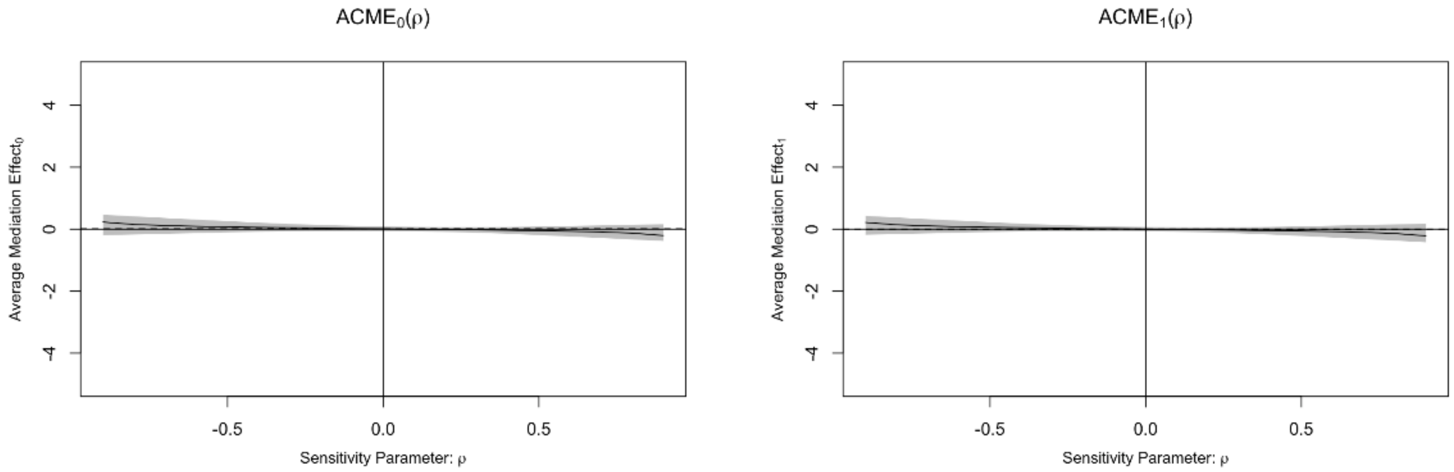

**Figure 1.2B**

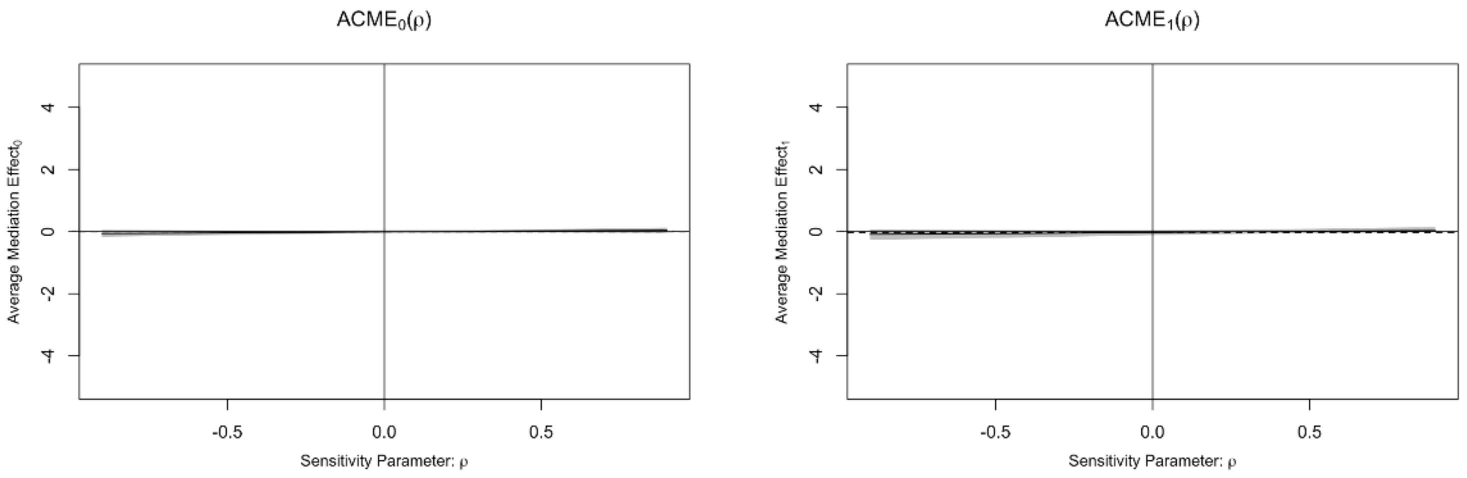


**Figure 1.2C**

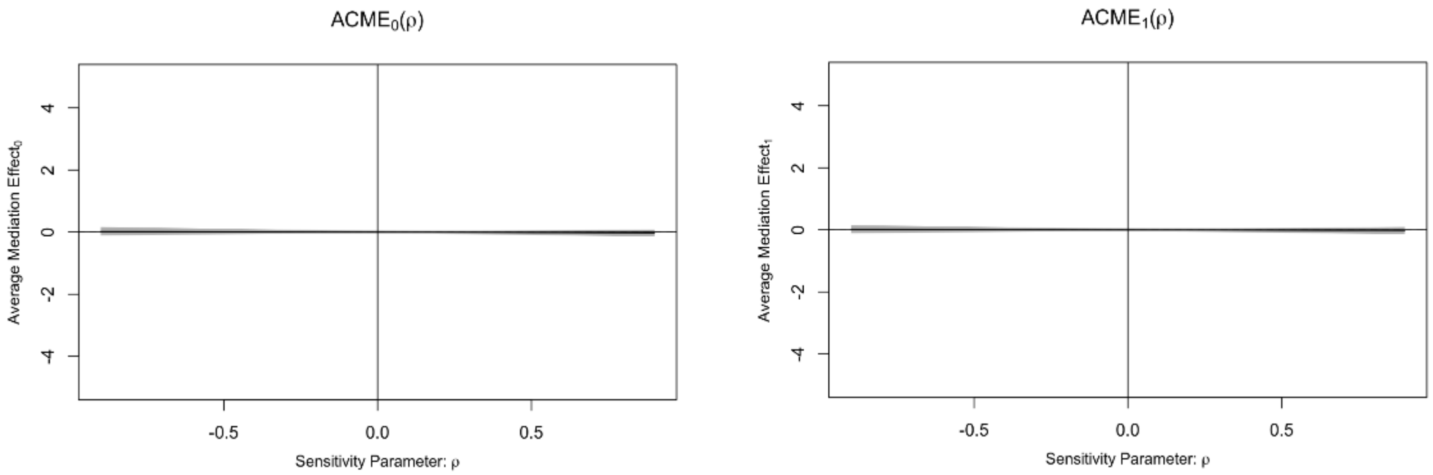


**Figure 1.2D**

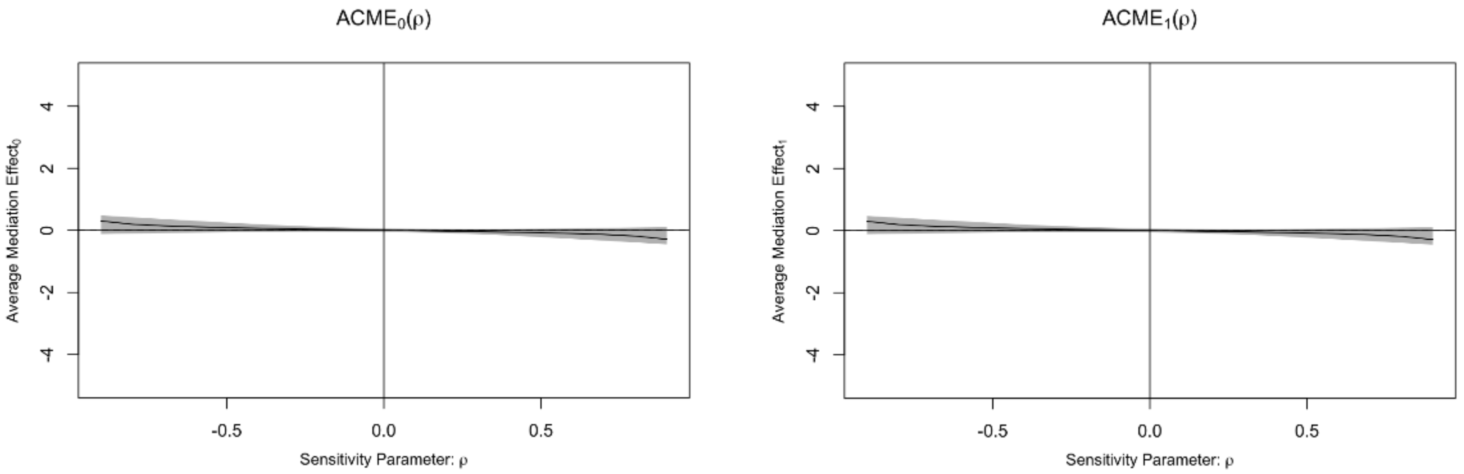

**Figure 1.3A**

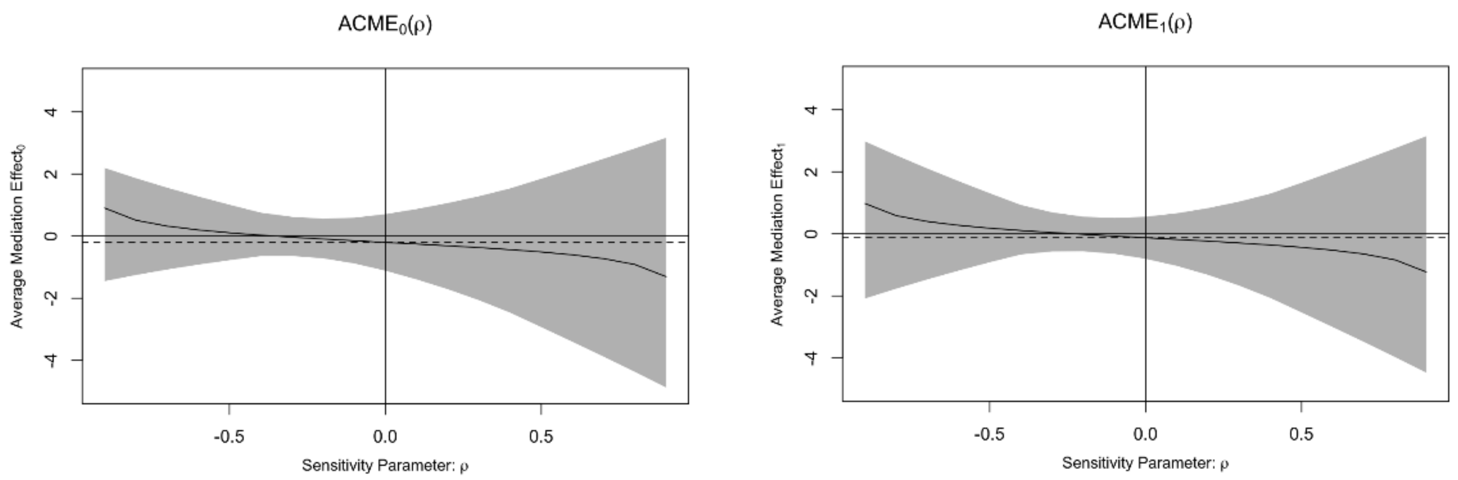


**Figure 1.3B**

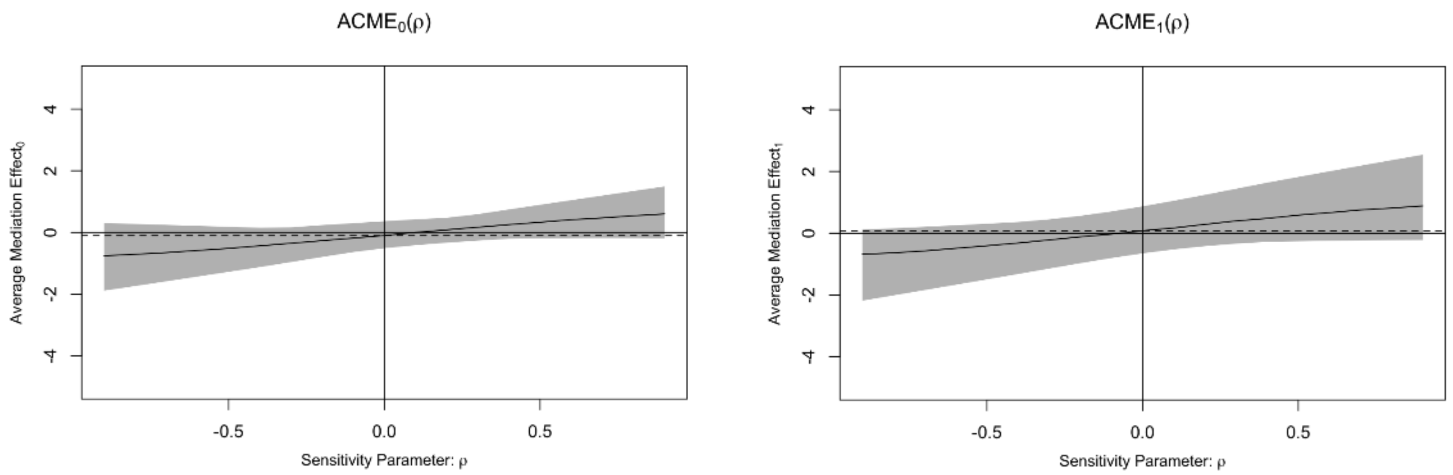


**Figure 1.3C**

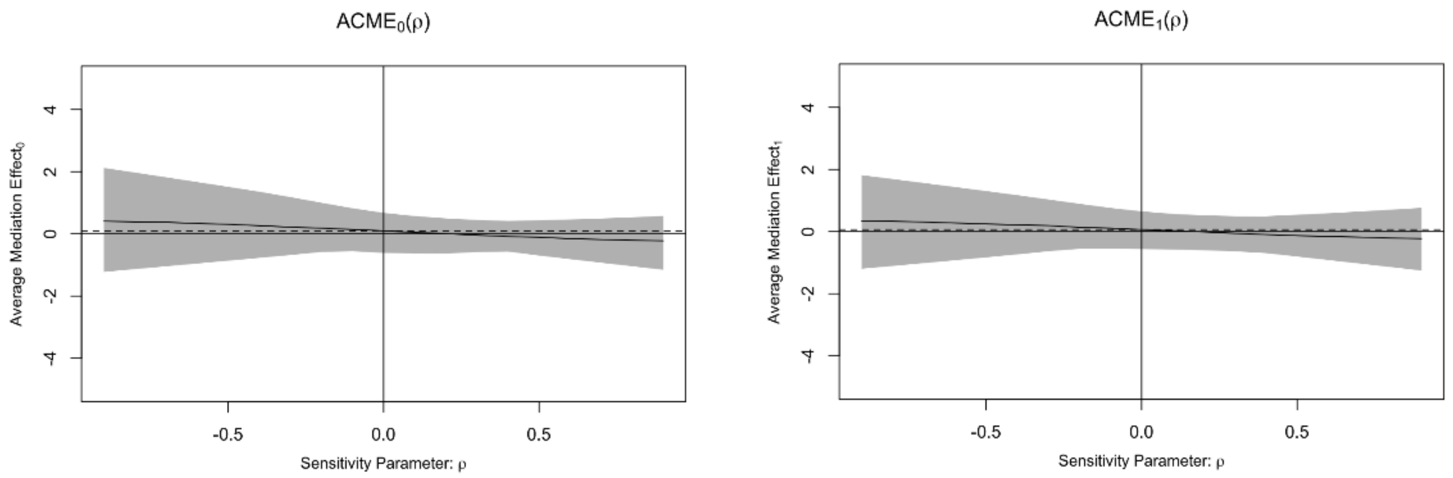

**Figure 1.3D**

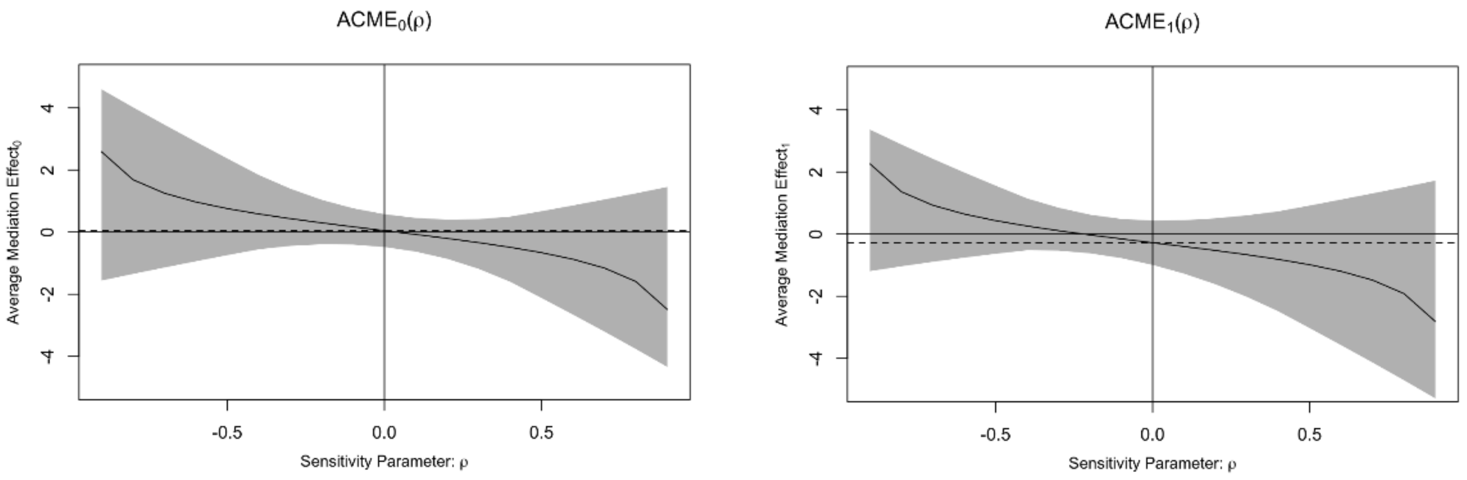


**Figure 1.4A**

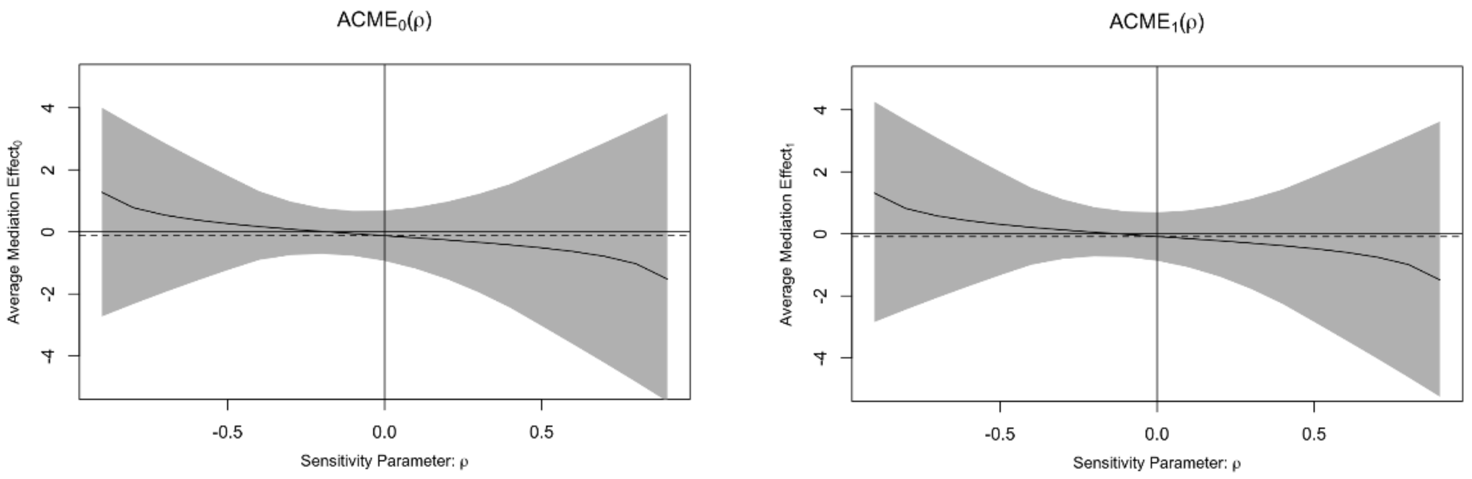


**Figure1.4B**

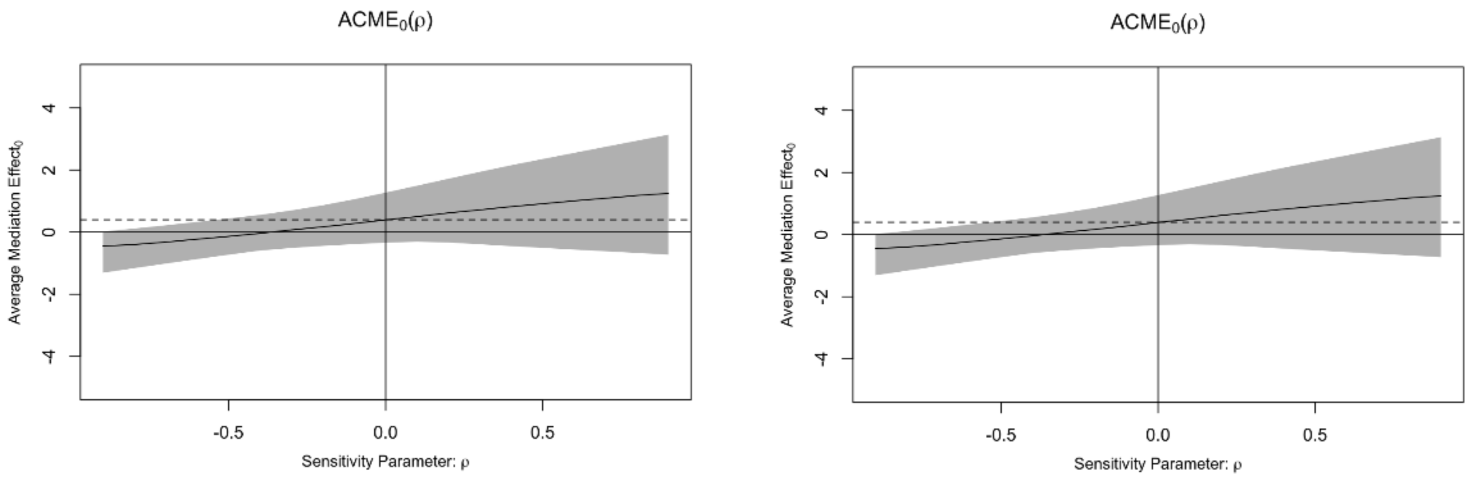

**Figure1.4C**

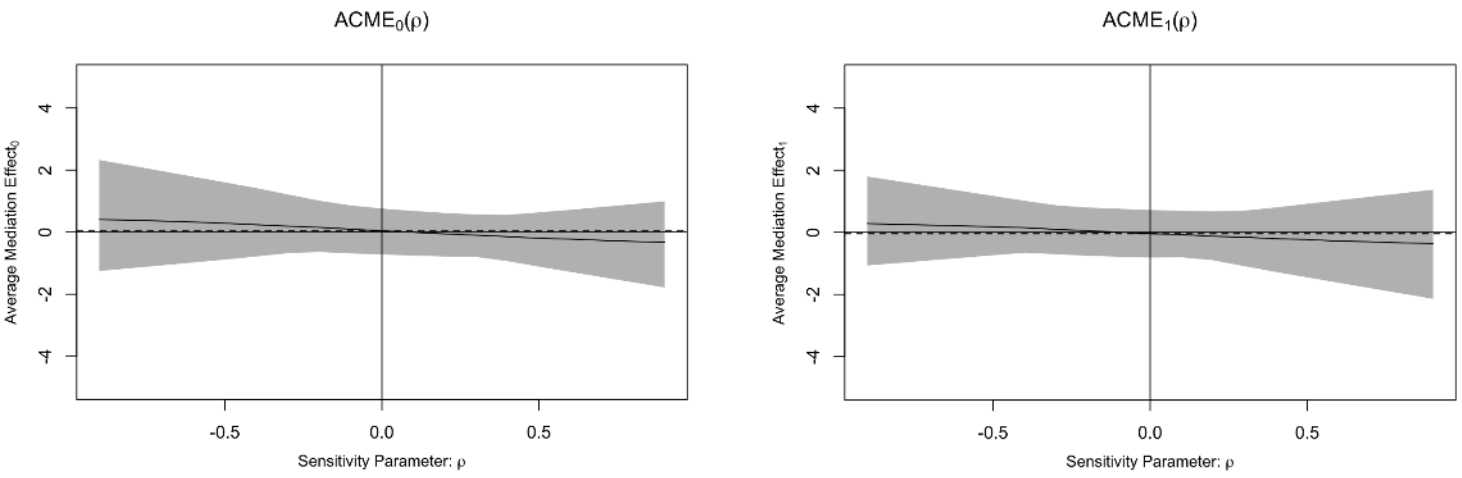


**Figure 1.4D**

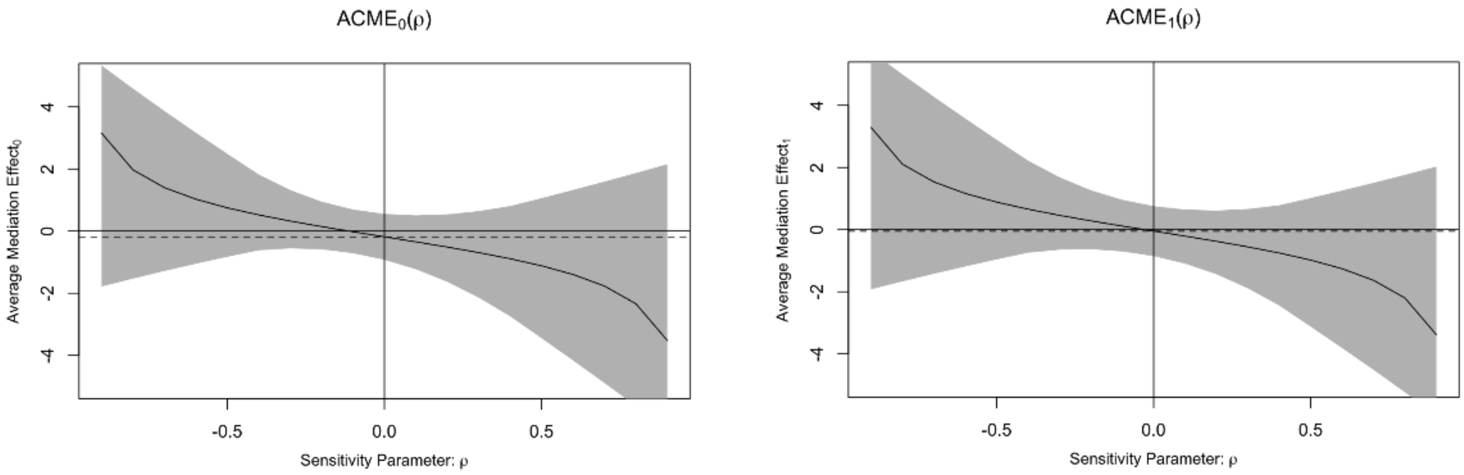
